# Temporal fingerprints of TMS-evoked potentials across thalamocortical circuits

**DOI:** 10.64898/2026.06.29.734769

**Authors:** Gabriel Hassan, Gianluca Gaglioti, Giulia Furregoni, Elena Focacci, Marta Porro, Letizia Bernardelli, Alessandra Calcagno, Marcello Massimini, Simone Sarasso, Mario Rosanova, Silvia Casarotto

## Abstract

**Background:** Electroencephalographic (EEG) potentials evoked by transcranial magnetic stimulation (TMS) offer a direct window into cortical dynamics. Yet, a systematic exploration of their morphological features, analogous to sensory-evoked potentials, is lacking, especially for stimulation outside the motor cortex.

**Aim:** To obtain region-specific properties of frontal, parietal and occipital networks from the time course of TMS-evoked potentials (TEPs).

**Materials and Methods:** We implemented and applied an automatic procedure to compute peak-to-peak amplitude, peak latency, and inter-peak interval of TEPs recorded from 40 neurotypical subjects stimulated over left occipital (n=25), parietal (n=25), and frontal (n=25) cortices.

**Results:** Occipital TEPs showed the largest peak-to-peak amplitude and longest latency of the first waveform component, independently of stimulation intensity and consistent with the recruitment of a large patch of densely interconnected neurons. Concerning later components, both latency and inter-peak interval systematically decreased along the posterior-to-anterior axis, reflecting progressively faster recurrent dynamics from the alpha-dominated occipital circuitry to the tightly coupled loops between frontal cortex and subcortical structures. Parietal TEPs showed intermediate amplitude and latency measures, consistent with the heterogeneous cytoarchitectonic and connectional organization of the superior parietal cortex.

**Conclusions:** Our findings suggest that TEP morphology is shaped by the distinct properties of the stimulated networks, with early amplitude reflecting the extent of local recruitment and later temporal features tracking the rhythm of recurrent activity. This work offers a mechanistically grounded and practically accessible approach, also released as a Python-based tool, that allows to characterize cortical reactivity across different brain-states and populations.

## 1. Introduction

The morphology of electroencephalographic (EEG) potentials — whether elicited by sensory stimulation or by direct cortical perturbation with transcranial magnetic stimulation (TMS) — reflects key aspects of the timing, strength, and organization of neural circuit responses. When stimulation occurs through peripheral pathways, the resulting sensory-evoked potentials (EPs) are classically described by the latency and amplitude of waveform components (Dawson, 1954), which are considered reproducible, clinically validated markers of neural pathway integrity (Chiappa and Ropper, 1982). A few paradigmatic clinical applications include the visual evoked potential (VEP) P100 in demyelinating disorders (Halliday et al., 1972), the somatosensory evoked potential (SEP) N20 as a predictor of poor outcome in post-anoxic coma (Cruccu et al., 2008; Logi et al., 2003), and auditory brainstem responses (ABR) in the diagnosis of retrocochlear pathologies (Starr and Achor, 1975). The interpretive power of these measures rests on a shared rationale: each component reflects a discrete, anatomically defined stage of neural processing. While well established, conventional EPs allow only limited inference, restricted to the specific sensory circuits engaged by peripheral stimulation.

The combination of TMS and EEG (TMS–EEG) overcomes this constraint by enabling direct perturbation of virtually any accessible cortical target, offering a unique window into local and distributed cortical reactivity (Atalay et al., 2025; Massimini et al., 2005; Russo et al., 2025; Ziemann et al., 2026). An analogous waveform-based approach to sensory EPs has been extensively developed for primary motor cortex (M1), where TMS-evoked potentials (TEPs) have been decomposed into canonical components (Beck et al., 2024; Farzan and Bortoletto, 2022; Premoli et al., 2014). Yet M1 represents a special case, in which TEPs are shaped by motor evoked potentials-guided target selection, by its peculiar cytoarchitecture — especially at the hand hotspot — and by potential confounds related to sensory reafference (Fecchio et al., 2017).

Non-motor TEPs can differentiate brain states and show promising clinical applications across psychiatric, neurodegenerative, and brain injury populations (Casarotto et al., 2019; Donati et al., 2023; Sarasso et al., 2020). Yet, unlike sensory EPs, they have not been systematically characterized through scalable, quantitative time-domain features. This gap is notable because cortical responses are known to carry region-dependent signatures shaped by cytoarchitecture, myeloarchitecture, and large-scale connectivity (Frauscher et al., 2018; Lysomiski et al., 2026; Momi et al., 2025; Parmigiani et al., 2022; Rosanova et al., 2009). Whether such regional specificity is systematically encoded in TEP morphology, and whether it can be captured with the rigor of standard EP analysis, remains an open question.

Here, we address this question by implementing and applying a standardized, semi-automatic, quantitative framework for characterizing TEP morphology across thalamocortical networks. We computed time-domain features — including peak-to-peak amplitude, latency, slope and inter-peak interval of waveform components — from 75 TMS-EEG sessions in 40 neurotypical subjects stimulated on the left occipital, parietal, and frontal cortices. These time-domain features significantly differentiate the TEPs across all targets and provide novel insight on the mechanisms governing circuit reactivity in different cortical areas. In addition, to facilitate further explorations and comparisons, we release the analysis pipeline as an open-source Python-based tool ([GITHUB link TBD], Figure 1).

**Figure 1.**
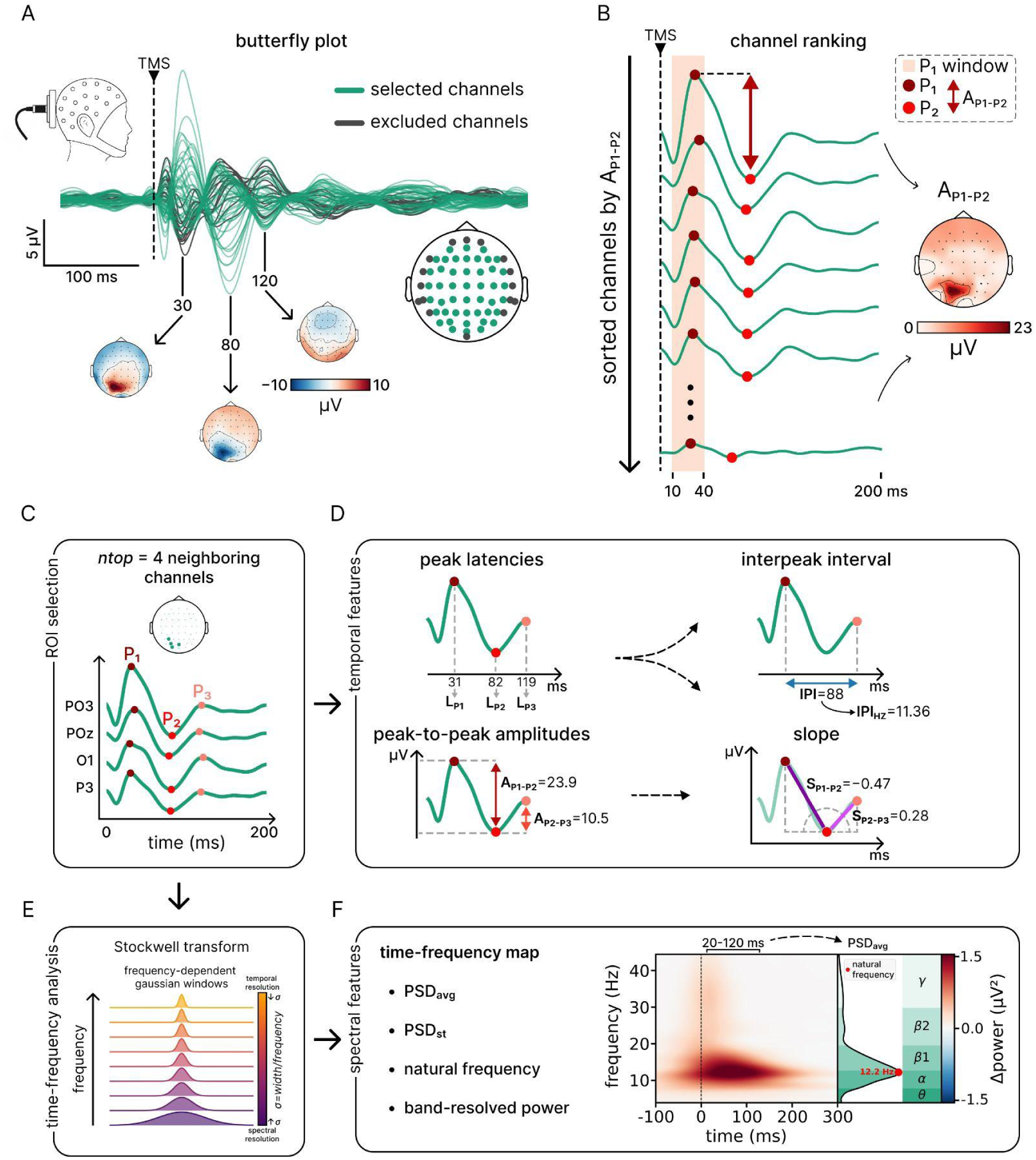
Data-driven procedure to compute TEP features in the time and frequency domains. **(A)** Butterfly plot of average TEPs of a representative subject during left occipital stimulation. Green traces and dots represent the 46 channels retained for data analysis. Lateral channels (in black) have been excluded because of low signal-to-noise ratio (see paragraph 2.5). **(B)** Single-channel TEPs sorted according to the peak to peak amplitude between P_1_ and P_2_ peaks (A_P1-P2_). P_1_ (dark-red dot) is detected within the time window between 10 and 40 ms (shaded area);P_2_ (red dot) is the following peak with opposite polarity. **(C)** An ROI of ntop = 4 channels was automatically selected from the channel with largest A_P1-P2_ plus 3 adjacent channels sorted by A_P1-P2_ and forming a connected cluster.P_3_ (light-red dot) is the peak following P_2_ with the same polarity as P_1_. **(D)** time-domain features automatically computed in each ROI channel: (i) *latency* of P_1_, P_2_, and P_3_ (L_P1_, L_P2_, L_P3_) in ms; (ii) *peak-to-peak amplitude* between P_1_ and P_2_ (A_P1-P2_) and between P_2_ and P_3_ (A_P2-P3_) in μV; (iii) *slope* of the components between P_1_ and P_2_ (S_P1-P2_) and between P_2_ and P_3_ (S_P2-P3_) in μV/ms as the ratio between the corresponding peak-to-peak amplitude and latency; (iv) *inter-peak interval* (IPI) as the time lag between P_1_ and P_3_ (=L_P3_-L_P1_) in ms and its reciprocal transformation in Hz (IPI_Hz_=1000/IPI). all measures were averaged across the 4 ROI channels before comparing stimulation sites. **(E)** Single-channel average TEPs belonging to the 4 ROI channels are analyzed in the time-frequency domain via Stockwell transform. **(F)** The power within the post-stimulus time window 20-120 ms is computed to derive the power spectral density of average TEPs (PSD_avg_ and its peak frequency represents the *natural frequency* (NF, red dot on the PSD_avg_). Band-resolved power is computed as the area subtended by the PSD_avg_ (green shading) within θ (4-7Hz), α (8-13Hz), β1 (13-20Hz), β2 (20-30Hz) and γ (30-45 Hz) bands.

## 2. Materials and Methods

### 2.1 Subjects

We collected 75 TMS-EEG sessions in 40 healthy adult subjects (mean age = 32.7 ± 9.8; 20 female). Enrolled subjects were screened for contraindications to TMS and did not declare any relevant medical, neurological or psychiatric disorder as well as use of substances potentially affecting brain functions. All the procedures conformed to the Declaration of Helsinki and were approved by the local ethics committee (Comitato Etico Milano Area 1; TMS-EEG Prot. n. 609/07/27/05/AP). Subjects signed a written informed consent before participation.

### 2.2 Experimental protocol

Single biphasic TMS pulses were delivered via a figure-of-8 coil using an individual MRI-based Navigated Brain Stimulation system (Nexstim Ltd., Finland), which also logged the induced maximum electric field (EF-max) on individual cortical targets; TMS-evoked responses were recorded with a 64-channel TMS-compatible BrainAmp DC amplifier at 5000 Hz (Brain Products GmbH, Germany). See Supplementary for more details.

During recordings, subjects were seated comfortably wearing in-ear earphones playing customized noise to mask the TMS click (Russo et al., 2022). Three TMS targets were visually identified on each subject’s 3D-rendered brain: the superior occipital gyrus (BA18/19), superior parietal gyrus (BA7), and superior frontal gyrus (BA6) of the left hemisphere.

Each session began by orienting the induced EF orthogonal to the targeted gyrus at an intensity corresponding to ∼100 V/m on the gyral crown (%MSO). Stimulation parameters (location, orientation, intensity) were then individually adjusted using a real-time TEP visualization tool (Casarotto et al., 2022) to minimize scalp muscle artifacts on single-trial data and to achieve an early (10–50 ms) peak-to-peak TEP amplitude >6 μV in average reference at the channel beneath the coil.

We recorded 25 TMS-EEG sessions per target across 40 subjects, yielding 75 TEP recordings overall. Each session included 150–250 stimuli with a jittered ISI of 2–2.3 s. Some subjects were stimulated at only 1 or 2 targets; 10 subjects completed all three, making 30/75 TEPs within-subject measurements.

### 2.3 Data analysis

EEG data were preprocessed following the pipeline described in Casarotto et al. (2024), including TMS pulse artifact removal (between −2 to +5ms), high-pass filtering at 1Hz, epoching (−600 to +600 ms), rejection of artifactual channels and trials by visual inspection, average re-referencing, ICA-based ocular and muscular artifacts removal, low-pass filtering at 45 Hz, and interpolation of bad channels. See Supplementary for more details.

TEP features in the time and frequency domain were computed from a subset of 46 EEG channels (Fig. 1A, green dots and traces). Excluded channels comprised the outermost lateral electrodes (Fp1, Fpz, Fp2, F7, F8, FT7, FT8, T7, T8, TP7, TP8, TP9, TP10, P7, P8, Iz; Fig. 1A, black dots and traces), which typically show a low signal-to-noise ratio due to their distance from the stimulated targets and their susceptibility to residual high-frequency artifacts deriving from facial and scalp muscle activity.

To compare the coordinates of the stimulation targets across subjects, individual MRIs underwent a cortical reconstruction process using the FreeSurfer software (Fischl, 2012) and were normalized to a standard brain template. This transformation was then applied to the 3D spatial coordinates of the EF-max to derive the corresponding stimulation coordinates in the Montreal Neurological Institute (MNI) reference space.

#### 2.3.1 Spatio-temporal TEP features

For each channel, we first searched for an evoked peak (P_1_) within a pre-defined time window between +10 and +40 ms after the pulse (Fig. 1B, shaded area). To automatically determine its polarity, we compared the TEP voltage at the midpoint of this time window (25 ms) with the average TEP voltage at its edges (i.e., at 10 and 40 ms): if midpoint voltage exceeded the edge average, the signal was concave and P_1_ was identified as a local maximum (*positive peak*); otherwise, the signal was convex and P_1_ was identified as a local minimum (*negative peak*). Then, we searched for two subsequent peaks: P_2_ as the first peak following P_1_ and having an opposite polarity, and P_3_ as the first peak following P_2_ and having the same polarity as P_1_. As an additional constraint, to avoid any possible distortion around the cutoff frequency introduced by low-pass filtering (see Data preprocessing paragraph), we set a minimum time lag of 14 ms between consecutive peaks, which corresponded to the half-period of a sinusoid at 35 Hz.

The following latency and amplitude features of average TEPs were computed for each channel (Fig. 1D): (i) absolute peak-to-peak amplitude between P_1_ and P_2_ (A_P1-P2_) and between P_2_ and P_3_ (A_P2-P3_) in µV; (ii) peak latencies of P_1_, P_2_ and P_3_ in ms (L_P1_, L_P2_, L_P3_); (iii) slope of the components between P_1_ and P_2_ (S_P1-P2_) and between P_2_ and P_3_ (S_P2-P3_) as the ratio between peak-to-peak amplitude and latency; (iv) interpeak interval (IPI) as the time lag between P_1_ and P_3_ in ms (i.e., L_P3_-L_P1_) and its convenient reciprocal transformation in Hz (IPI_Hz_=1000/IPI). IPI and IPI_Hz_ correspond, respectively, to the period and to the frequency of the first oscillation evoked by TMS.

To focus subsequent analyses on electrodes most sensitive to local cortical reactivity, channels were ranked in descending order according to A_P1-P2_. We defined a region-of-interest (ROI) of n channels (*ntop* channels) with the highest A_P1–P2_ amplitude, constrained to form a spatially connected cluster (Fig. 1C). Specifically, the ROI selection started from the highest-ranked channel, then expanded by iteratively adding the neighboring channels with the next highest A_P1–P2_ values, ensuring spatial contiguity at each step. This ROI of neighboring channels was then used in second-level analyses to compare local TEP features among stimulation targets.

#### 2.3.2 Spectral TEP features

Spectral TEP features between 4 and 45 Hz were estimated by applying the Stockwell transform, a time-frequency decomposition method that combines Short Time Fourier Transform with signal filtering by frequency-dependent Gaussian windows (Stockwell et al., 1996). The width of the Gaussian window is used to balance temporal and frequency resolution: values below 1 favor temporal resolution, while values above 1 favor frequency resolution. In the following analyses, we set width = 0.7 unless otherwise specified. For each channel, we computed time-frequency power spectra of average TEPs (TF_avg_) and we applied a baseline correction by subtracting the average pre-stimulus power (from −500 to −100 ms) from each frequency bin. Then, we computed the power spectral density (PSD_avg_) evoked by TMS by summing baseline-corrected power within a selected post-stimulus time window between 20 and 120 ms. Finally, we estimated the natural frequency (NF) as the frequency bin corresponding to the maximum of the power spectral density (Fig. 1F). In parallel, time-frequency power spectra were also computed from single-trial EEG responses for each channel and averaged across trials; then they were subjected to the same analysis pipeline to derive a trial-based estimate of baseline-corrected PSD (PSD_ST_) and the corresponding natural frequency (NF_ST_).

#### 2.3.3 Statistical analysis

All statistical analyses were performed in Python (SciPy, scikit-posthocs, statsmodels). Non-parametric methods were used throughout due to violations of normality (Shapiro-Wilk) and/or homoscedasticity (Levene). Between-site comparisons relied on Kruskal-Wallis tests followed by Dunn’s post-hoc tests with Holm-Bonferroni correction. Within-subject analyses (n = 10, all three targets) used Friedman tests with Wilcoxon signed-rank post-hoc tests and Holm-Bonferroni correction. Pairwise feature correlations were assessed with Pearson’s r, with p-values adjusted via Benjamini-Hochberg false discovery rate correction; the relationship between inter-peak interval and natural frequency was further characterized by ordinary least squares regression. Data are presented as mean ± SD; box plots depict median, interquartile range, and 5th–95th percentiles.

## 3. Results

We investigated the EEG responses to TMS of three distinct cortical targets over the left hemisphere. We characterized the intrinsic properties of these different cortical circuits by applying a semi-automatic analysis pipeline that computes synthetic spatial, temporal and spectral features of TEPs. The centroid of each stimulation target, computed as the median EF-max coordinates in MNI reference space across subjects, was located on the superior frontal gyrus (*x* = −16.7, *y* = 9.8, *z* = 64.6 mm), superior parietal gyrus (*x* = −21.5, *y* = −67.7, *z* = 60.7 mm) and superior occipital gyrus (*x* = −13.4, *y* = −88.3, *z* = 30.2 mm). On average, we retained 200.6 ± 45.7 trials for each TMS-EEG session, without significant (one-way ANOVA; p>0.05) differences among stimulation targets (min-max number of trials: 112-282 for frontal, 82-248 for parietal and 103-289 for occipital targets). The number of bad channels ranged from 0 (55 out of 75 TMS-EEG sessions) to 3. The signal-to-noise ratio, computed as the ratio of the mean absolute amplitude between each of two post-stimulus time windows (from 10 to 50 ms and from 10 to 120 ms, respectively) and a pre-stimulus time window (from −400 to −10 ms), was above 7 and 4.4 respectively for all sessions (Supplementary Fig. S1).

### 3.1 Topography of early TEP amplitude across stimulation targets

To assess the spatial distribution of early TEP amplitude, we identified for each participant and target a progressively larger set of neighboring channels (*ntop* parameter; Fig.1C) starting from the one with the highest A_P1-P2_ value. For each *ntop* value, we then calculated the maximum percentage of participants sharing at least one channel within their respective ROIs (Fig. 2A). We found that, with *ntop* = 4, all participants shared electrode FC1 following frontal stimulation, whereas electrode PO3 was common to all participants following occipital stimulation (Fig. 2B), indicating a high degree of consistency in TEP topographies across individuals. A different pattern emerged for parietal stimulation, where only 56% of participants shared a common channel at ntop = 4 (Fig. 2B). Nevertheless, this variability was structured rather than random: electrode C1 was shared by 12 participants, electrode P1 by 11 participants, and both C1 and P1 by the remaining 2 participants, suggesting that the largest A_P1-P2_ values were consistently centered around either C1 or P1, with an approximately equal distribution across individuals (Fig. 2B and Supplementary Fig. S2A). Importantly, these findings could not be explained by differences in stimulation targeting, as individual EF-max locations clustered over the superior frontal, parietal, and occipital gyri in accordance with the experimental protocol (Fig. 2A), and the spatial dispersion of parietal targets was not significantly greater than that observed for frontal or occipital targets (Supplementary Fig. S2B).

**Figure 2.**
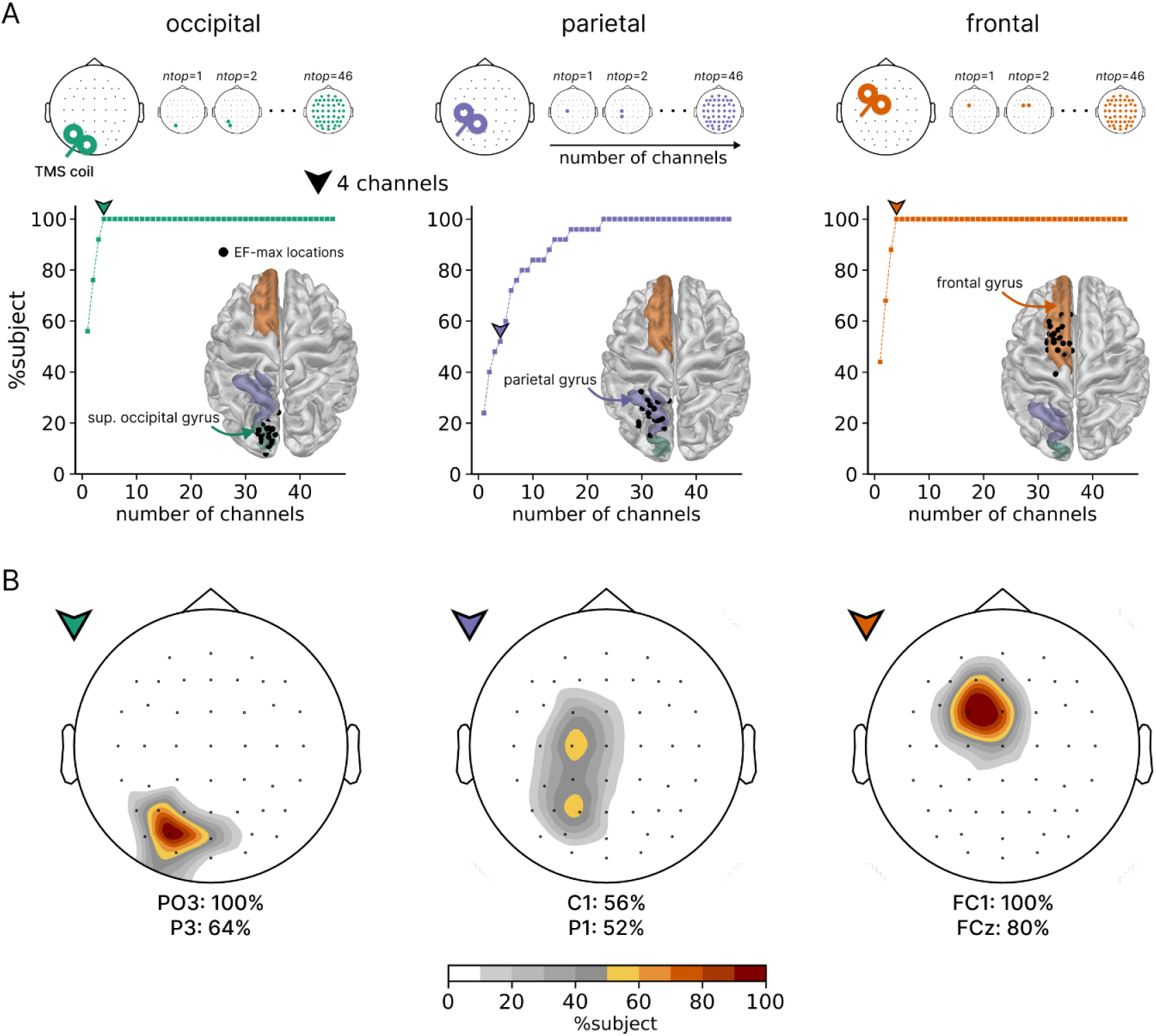
Spatial distribution of TEPs amplitude and of stimulation coordinates. **(A)** Largest percentage of subjects with at least one channel in common among the individual ROI channels automatically identified based on largest A_P1-P2_, by varying the *ntop* parameter. The percentage of subjects corresponding to *ntop* = 4 is highlighted by an arrow tip. On each template brain, black dots represent individual EF-max locations in MNI coordinates whereas color-shaded areas depict the superior occipital, parietal and frontal gyri according to the Destrieux atlas (Destrieux et al., 2010). **(B)** Topographic display of the percentage of subjects sharing the same channel among the ROI channels automatically identified with *ntop* = 4 (arrow tip). In all panels, green, violet and orange colors indicate occipital, parietal and frontal targets, respectively.

### 3.2 Temporal profile across cortical sites

The stimulation of different cortical targets elicited EEG responses with distinct temporal dynamics, as suggested by the qualitative differences in TEP morphology observed in representative butterfly plots (Fig. 3A). To quantify these differences at the group level, we extracted a set of temporal TEP features from signals averaged across the *ntop* = 4 ROI channels in each participant (Fig. 1C-D). Inspection of grand-average TEP waveforms as well as the latency of evoked components (P_1_, P_2_ and P_3_ peaks (Fig. 3B) revealed a progressive temporal shift across stimulation sites, with occipital stimulation eliciting slower oscillatory responses than parietal and frontal stimulation. Consistent with this observation, L_P1_ was significantly longer for occipital than frontal TEPs (Fig. 3C), whereas L_P2_ was significantly prolonged for occipital TEPs relative to both parietal and frontal TEPs (Fig. 3D). Notably, L_P3_ (Fig. 3E) as well as IPI (Fig. 3F) decreased progressively from occipital to parietal to frontal stimulation, with significant differences across all pairwise comparisons. In addition to these latency effects, we found significantly larger A_P1-P2_ following occipital stimulation as compared to either parietal or frontal stimulation (Fig. 3G, Fig. S3A). This finding cannot be explained by stimulation intensity, as expressed either by %MSO or by the induced EF-max, which were significantly higher in parietal compared to both frontal and occipital targets. (Fig. S3B-C). Conversely, A_P2-P3_ did not differ across stimulation sites (Fig. 3H). Finally, slope measures, which are likely to be affected by both peak amplitude and latency, showed that absolute S_P1-P2_ was comparable across sites (Fig. 3I), whereas absolute S_P2-P3_ significantly discriminated among the three cortical targets (Fig. 3J).

**Figure 3.**
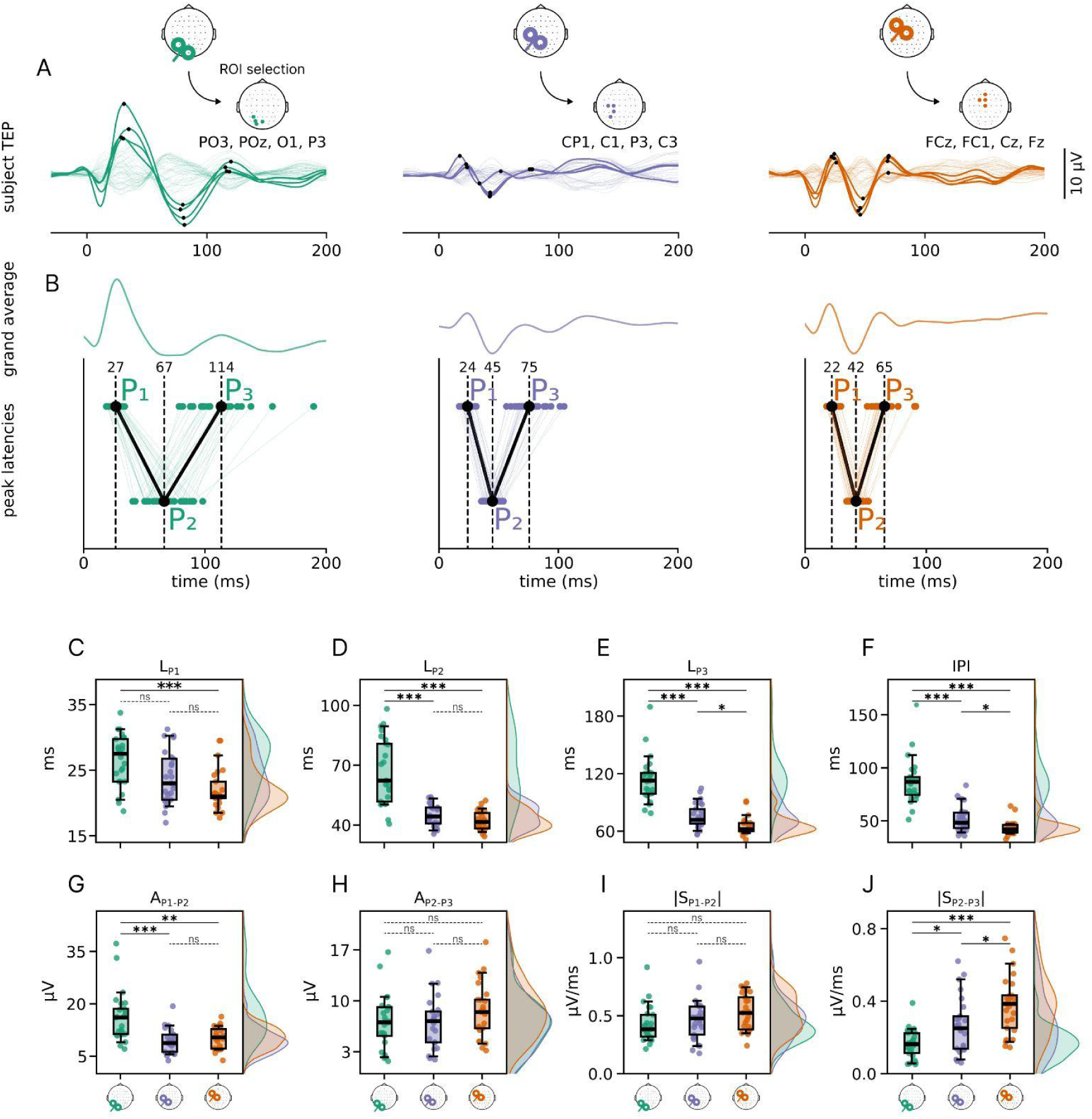
Morphological features of TEPs across stimulation sites. **(A)** Butterfly plot of TEPs elicited by stimulating the occipital (green), parietal (violet) and frontal (orange) targets in a representative subject. Bold traces and dots correspond to the ROI channels automatically identified based on largest A_P1-P2_ and setting ntop = 4. Black dots on the traces mark the automatically detected P1, P2, and P3 peaks. (B’) ROI-TEPs, obtained by averaging TEPs across the 4 ROI channels for each stimulation site, averaged across all subjects. **(B)** Colored dots represent single-subject latency of P1, P2 and P3 peaks averaged across the 4 ROI channels for each stimulation site. Black dots indicate the mean of L_P1_, L_P2_ and L_P3_ across subjects. Colored and black solid lines connect the corresponding dots in temporal sequence. **(C–J)** Group-level distributions of L_P1_ (C), L_P2_ (D), L_P3_ (E), IPI_Hz_ (F), A_P1-P2_ (G), A_P2-P3_ (H), |S_P1-P2_| (I) and |S_P2-P3_| (J) are depicted as boxplots (median, 25th and 75th percentile and 5th and 95th) and as density plots for occipital (green), parietal (violet) and frontal (orange) targets. Reported P-values were obtained by applying the Kruskal-Wallis test separately to each measure, followed by pairwise Dunn’s post-hoc tests with Holm-Bonferroni correction. In the figure, significance is indicated as * for p < 0.05, ** for p < 0.01, *** for p < 0.001, and ns for non-significant comparisons (p ≥ 0.05).

Analyses restricted to the subset of 10 participants who underwent all stimulation conditions yielded comparable results, confirming the robustness of the observed differences across cortical targets in a within-subject design (Supplementary Fig. S4A-H).

### 3.3 Spectral features across cortical sites

Time-frequency analysis of TEPs revealed clear spectral differences across stimulation sites (Fig. 4A). The PSD_avg_, obtained by summing baseline-corrected TEP power between 20 and 120 ms post-stimulus, exhibited distinct peaks depending on the stimulated cortical region (Fig. 4B). Specifically, the maximum of grandaverage PSD_avg_ occurred at around 11.7, 20.0, and 24.9 Hz for occipital, parietal, and frontal stimulation, respectively. Consistent with these observations , the natural frequency (NF) increased systematically along the posterior-to-anterior axis (Fig. 4C), reaching median values of 12.2Hz (IQR = 2.4Hz), 20.5 Hz (IQR = 2.9Hz), and 24.9Hz (IQR = 3.4Hz) for occipital, parietal, and frontal TEPs, respectively. All pairwise comparisons were statistically significant.

**Figure 4.**
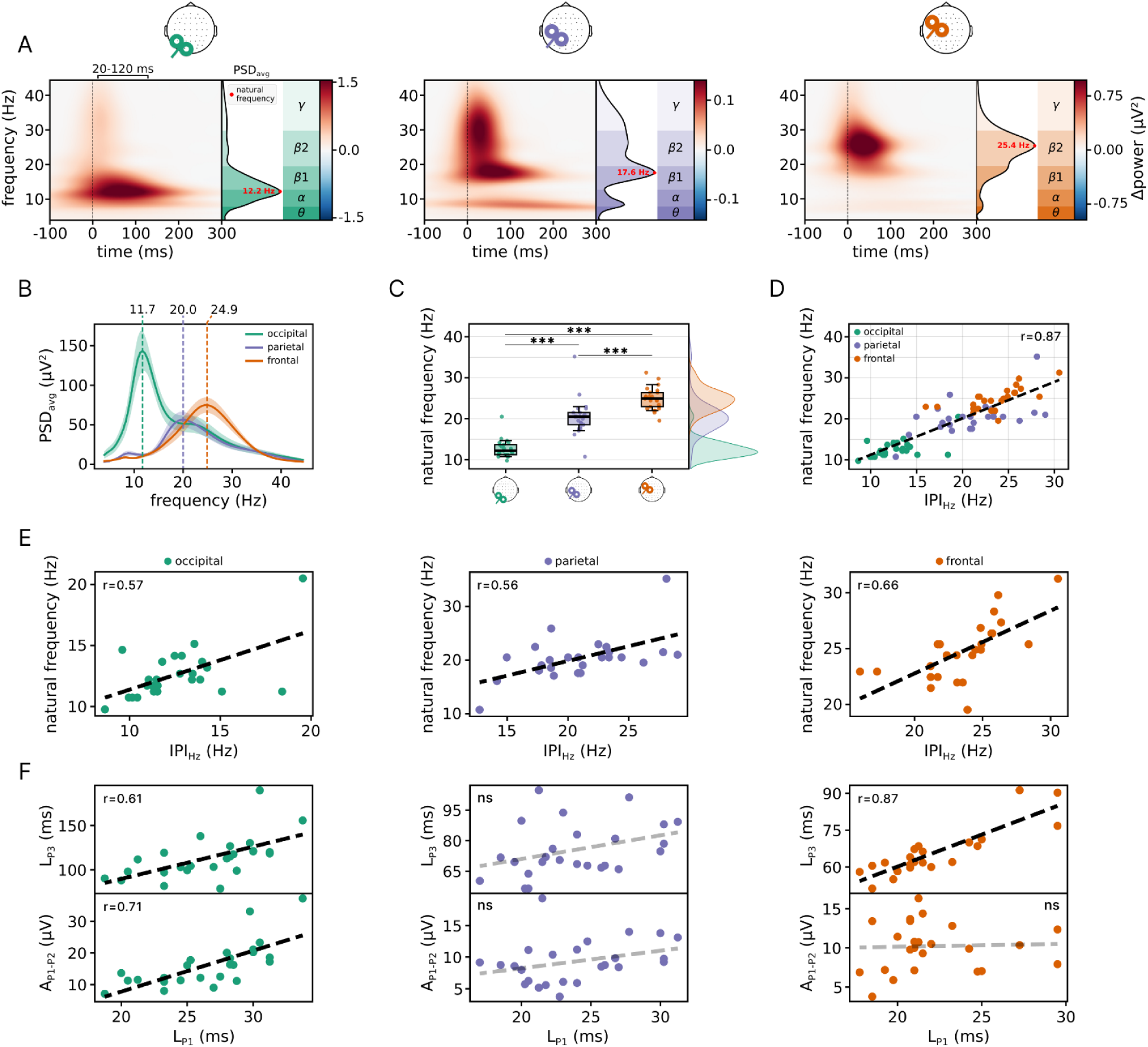
Time-frequency analysis of TEPs reveals stimulation-specific natural frequencies (NF) and relationships with temporal features. **(A)** Time–frequency power representations of a representative subject following occipital (left), parietal (middle), and frontal (right) stimulation applied on the four automatically selected channels. **(B)** Grand-average of the evoked power spectra across subjects for the three stimulation conditions. Shaded areas indicate the standard error of the mean (SEM).\ **(C)** Box plots of NF across subjects for each stimulation site (occipital: green, parietal: violet, frontal: orange), with overlaid subject-level distributions. Post-hoc pairwise comparisons (Dunn test), corrected using the Holm–Bonferroni method, were significant for all contrasts ( p < 0.001) and are indicated in the figure by ***. Density plots to the right of the box plot summarize the NF distribution across subjects for each stimulation site. **(D)** Linear correlation (r = 0.84, p = 4×10^-21^) between NF and IPI_Hz,_ across all conditions. The IPI_Hz_ converts the seconds-based interpeak intervals to Hz. . **(E)** Linear correlations between NF and IPI_Hz_, shown separately for occipital (r = 0.57, p = 0.0029), parietal (r = 0.56, p = 0.0038), and frontal (r = 0.66, p = 0.00037) stimulation. **(F)** Linear correlations between relevant TEP features for each stimulation site. Top row: P_1_ versus P_3_ latency (L_P1_ vs L_P3_; occipital: r = 0.61, p = 0.0013; frontal: r = 0.87, p = 2×10^-8^); bottom row: P_1_ latency versus peak-to-peak amplitude (L_P1_ vs A_P1-P2_; occipital: r = 0.71, p = 6×10^-5^). Significant correlations are shown with black dashed regression lines; non-significant (ns) associations are shown in gray.

As expected from their definitions (see Methods, sections 2.5.1–2.5.2), NF and IPI_Hz_ showed a strong association, as both metrics capture the dominant oscillatory properties of the evoked response through complementary approaches. Indeed, a significant linear relationship between NF and IPI_Hz_ was observed when IPI_Hz_ was averaged across the 4 ROI channels (r = 0.87, p = 7×10⁻²⁴; Fig. 4D) as well as when it was computed from the single channel exhibiting the largest A_P1-P2_ (r = 0.87, p = 2×10⁻²⁴). These findings indicate that IPI_Hz_ provides a reliable estimate of NF directly from the TEP waveform, making it potentially suitable for real-time applications during data acquisition. Similar results were obtained when NF was estimated from single-trial PSD_st_ (Supplementary Fig. S5); also, within-subject analyses confirmed both the posterior-to-anterior NF gradient and the NF-IPI_Hz_ correlation (Supplementary Fig. S4E–F). Furthermore, systematically varying the NF analysis parameters (*ntop*, Gaussian window width, and post-stimulus time window) showed that both results held across all combinations tested, confirming that they are not a product of any particular parameter choice (Supplementary Fig. S6).

Finally, we examined the relationships among relevant features for each stimulation site (Fig. 4 E-F). IPI_Hz_ and NF were significantly correlated within each stimulation site, further consolidating their shared sensitivity to the circuit-level oscillatory properties of the evoked response. Additional site-specific associations also emerged. In particular, L_P3_ was positively correlated with L_P1_ across stimulation sites, suggesting that the latency of later TEP components is partly constrained by the timing of the initial response. Moreover, A_P1-P2_ was positively associated with L_P1_ exclusively for occipital stimulation. Collectively, these results indicate that each stimulation site is characterized by a distinct profile of temporal and spectral feature interactions, reflecting differences in the intrinsic dynamical properties of the underlying cortical circuits.

## 4. Discussion

In this study, we show that time-domain features of TMS-evoked potentials represent robust and reproducible fingerprints of cortical reactivity across three stimulation sites - occipital, parietal, and frontal - each embedded in a distinct thalamocortical network and characterized by a unique set of intrinsic circuit properties. Amplitude, latency, slope, and inter-peak interval measures, automatically computed from region-of-interest channels selected according to a data-driven procedure from evoked responses with a high SNR, reveal consistent site-dependent differences in TEP morphology and time course. This approach extends the identification of canonical features routinely used for sensory-EPs to cortical potentials evoked in areas beyond the primary motor cortex.

Focusing on the earliest component, A_P1-P2_ was significantly larger following occipital stimulation, a reproducible effect not explained by systematic changes in stimulation intensity or induced electric field strength. Moreover, occipital TEPs also exhibited a significantly longer L_P1_ compared to frontal TEPs, whereas no significant difference was observed relative to parietal stimulation. Together, these findings suggest that the earliest TEP component primarily reflects intrinsic circuit properties rather than a mere dose effect. One possible interpretation is that A_P1–P2_ and L_P1_ index the gain, synchrony, and temporal extent of local cortical recruitment immediately following the pulse, namely the efficiency with which the induced electric field engages a neuronal population capable of generating a coherent macroscopic response. Within this framework, unlike frontal and parietal cortices, the visual cortex is characterized by dense, spatially structured horizontal connections linking functionally related neuronal populations over millimeter scales (Gilbert and Wiesel, 1989; Stettler et al., 2002; Ts’o et al., 1990). Such lateral connectivity may facilitate both rapid synchronization and recruitment of a larger patch of interconnected neurons following focal perturbation, providing a plausible substrate for the larger amplitude and slightly prolonged latency of the early occipital response observed at the scalp level.

Beyond the earliest component, the latency of later peaks revealed progressively clearer differences across stimulation sites, outlining the electrophysiological fingerprint of the underlying circuits.

The evoked frontal pattern of activity is particularly informative in this respect. Compared to occipital and parietal stimulation, frontal TEPs were characterized by a shorter L_P3_ and by a tighter relationship between the latency of successive components, such that L_P3_ could be reliably predicted from earlier peak latencies. This finding suggests a more deterministic, chain-like temporal organization of the response, in which the timing of later components is strongly constrained by the timing of preceding ones. Such a pattern is compatible with sequential recruitment along recurrent pathways endowed with relatively stable conduction delays.

One possible substrate for these dynamics is the extensive network of cortico-striato-thalamo-cortical loops associated with frontal associative regions. Converging evidence from neuroimaging, TMS, intracranial recordings, and anatomical studies indicates that frontal stimulation engages distributed circuits involving the basal ganglia and thalamus in addition to local cortical networks (Bestmann et al., 2004; Denslow et al., 2005; Peters et al., 2020; Solomon et al., 2024). Unlike cortico-cortical interactions, which may propagate through multiple alternative routes of variable length and synaptic composition (Markov et al., 2014), these recurrent subcortical loops are organized along relatively stereotyped anatomical pathways (Alexander et al., 1986; McHaffie et al., 2005). Information traverses a limited number of relay stations connected by heavily myelinated fiber bundles and characterized by robust topographic organization (Jones, 2001). As a result, signal transmission through these circuits is expected to exhibit comparatively stable conduction delays and reduced temporal variability, favoring the emergence of reproducible sequences of activation following perturbation (Cona et al., 2011). Although the present data cannot dissociate the relative contribution of cortical and subcortical generators, the observed temporal coupling is consistent with the idea that frontal associative networks support faster and more structured re-entrant dynamics than other cortical circuits. In this framework, frontal TEPs may reflect the rapid engagement of recurrent cortico-subcortical loops whose stereotyped architecture imposes strong temporal constraints on the unfolding of the evoked response.

At the opposite end of the spectrum, occipital responses appear to combine strong early gain with slower subsequent evolution. The larger A_P1-P2_, together with prolonged L_P2_, L_P3_, and inter-peak intervals, points to a regime in which strong initial synchronization is followed by relatively slow oscillatory reverberations. This temporal profile is consistent with the dominant role of alpha-frequency dynamics in posterior cortices and with evidence linking alpha oscillations to visual cortical excitability and thalamo-cortical interactions (Lőrincz et al., 2009; Romei et al., 2008). Notably, these macroscopic observations have plausible cellular correlates. Single-unit recordings in the cat visual cortex offer a direct cellular parallel: a TMS pulse evokes a strong early facilitation at intervals of ∼100 ms followed by a prolonged period of reduced firing lasting several seconds (Moliadze et al., 2003). Such findings suggest that the combination of high early gain and slower temporal evolution observed in occipital TEPs may reflect fundamental properties of visual cortico-thalamo-cortical circuits, spanning multiple spatial scales from local neuronal populations to large-scale electrophysiological responses (Reinhold et al., 2015).

The parietal cortex occupied an intermediate position between occipital and frontal cortices across both temporal and spectral measures, while also showing distinctive waveform features and greater inter-individual variability in their spatial distribution. This pattern is consistent with the known heterogeneity of the posterior parietal cortex, a highly associative region comprising multiple cytoarchitectonic subdivisions with distinct functional roles and connectivity profiles (Thomas Yeo et al., 2011; van den Heuvel and Sporns, 2013). Unlike primary sensory or frontal associative regions, the posterior parietal cortex is embedded within several partially overlapping large-scale networks and serves as a major hub for integrating sensory, motor, and cognitive information (Andersen and Cui, 2009; Rolls et al., 2022). Such architectural diversity may naturally give rise to intermediate dynamical properties and to the greater variability observed across individuals in the spatial organization of the evoked response. This heterogeneity may also have practical consequences for TMS-EEG measurements at the parietal target. The coexistence of structurally and functionally distinct neuronal populations within the stimulated volume may reduce the initial synchronization of evoked responses, lowering the effective gain of the early cortical response and requiring higher stimulation intensities to achieve an adequate signal-to-noise ratio. In addition, TMS effects are highly sensitive to local cortical geometry and coil orientation (Janssen et al., 2015; Opitz et al., 2011). Within a heterogeneous associative region such as the posterior parietal cortex, even subtle variations in the exact location of the induced electric field may shift recruitment toward partially distinct circuits, preferentially engaging different neuronal elements — including axons, inhibitory interneurons, or pyramidal neurons — each characterized by distinct electrophysiological response profiles (Mueller et al., 2014). As a consequence, small differences in stimulation targeting may have a greater impact on the resulting TEPs than in the more functionally homogeneous occipital or frontal targets. Together, these factors likely account for both the higher intensity thresholds and the increased variability in early TEP topography observed at the parietal compared to the occipital and frontal targets.

The observed gradient in temporal dynamics closely parallels the typical posterior-to-anterior increase in dominant TMS-evoked frequencies described by Rosanova et al. (2009). In our dataset, natural frequency increased systematically from occipital to frontal cortex and was tightly mirrored by the reciprocal time-domain measure IPI_Hz_. Despite being derived through fundamentally different approaches, both measures appeared to capture the same underlying dynamical property of the evoked response. This finding suggests that the duration of the first oscillatory cycle provides a simple temporal estimate of the dominant frequency at which the stimulated circuit reverberates following perturbation. More broadly, the close relationship between NF and IPI_Hz_ supports the idea that waveform morphology and spectral organization represent complementary descriptions of a common circuit-level process. In this framework, the inter-peak interval emerges as a compact synthetic index linking the temporal structure of TEPs to their oscillatory content, extending previous efforts to derive physiologically meaningful measures from TMS-evoked responses (Casali et al., 2010). From a practical perspective, IPI may provide a convenient surrogate of natural frequency in experimental and clinical settings requiring rapid online characterization of cortical dynamics.

These temporal features are not simple proxies for spectral measures, but are also well positioned to capture state-dependent modifications of cortical reactivity. For example, during NREM sleep and general anesthesia, TMS evokes a preserved or enhanced early response followed by a rapid collapse into a large, stereotyped slow wave (Ferrarelli et al., 2010; Massimini et al., 2005; Sarasso et al., 2014). Such sleep-like dynamics would be expected to manifest as a modulation of both amplitude and latency measures, consistent with a bistability-driven regime in which a strong transient activation is followed by a cortical OFF-period disrupting fast recurrent interactions (Pigorini et al., 2015; Rosanova et al., 2018). Similar patterns have been reported in disorders of consciousness and in the perilesional cortex after brain injury as well as in psychiatric conditions characterized by reduced fast oscillatory activity (Ferrarelli et al., 2012; Meneghini et al., 2025; Rosanova et al., 2018; Sarasso et al., 2020). Unlike spectral measures — which summarize the activity across the entire post-stimulus window — discrete peak latencies and amplitudes probe distinct moments of the evoked response, offering a more granular and complementary view of circuit-level dynamics. Characterizing these features may therefore prove physiologically informative, even if establishing normative values and standard ranges will ultimately require validation in larger datasets spanning different cortical targets, age groups and populations. Used in conjunction with spectral measures, this approach may enable a more precise dissociation of the circuit mechanisms underlying changes in cortical reactivity across brain states and disorders.

Another important implication of the present work is methodological. Outside the primary motor cortex, TEP morphology has often been described qualitatively or with heterogeneous analysis choices, limiting reproducibility and making comparisons across studies difficult (Farzan and Bortoletto, 2022). Here, we show that a semi-automatic workflow based on subject-specific yet standardized channel selection can provide a principled way to extract comparable time-domain features from non-motor TEPs. The specificity of these measures to recapitulate known spectral gradients further supports the view that waveform morphology contains structured information that can be systematically quantified and interpreted.

In conclusion, TEPs can be summarized by a set of simple, mechanistically interpretable features that capture distinct aspects of cortical reactivity across different networks. Early peak-to-peak amplitude indexes the gain of local recruitment, whereas later latencies and inter-peak intervals reflect the temporal scale of recurrent dynamics. This dissociation reveals a principled organization across cortical systems and provides a compact framework for characterizing thalamo-cortical function in both basic and clinical contexts. Finally, by extending the time-domain framework of conventional sensory evoked potentials to direct cortical perturbation, these results provide a bridge between classical electrophysiological approaches and network-level probing of brain intrinsic properties.

## CRediT authorship contribution statement

**Gabriel Hassan**: Writing – review & editing, Writing – original draft, Methodology, Formal analysis, Investigation, Data curation, Conceptualization. **Gianluca Gaglioti**: Writing – review & editing, Writing – original draft, Software, Visualization, Methodology, Formal analysis, Conceptualization. **Giulia Furregoni**: Investigation, Writing – review & editing. **Elena Focacci**: Investigation, Writing – review & editing. **Letizia Bernardelli**: Investigation, Writing – review & editing. **Marta Porro**: Investigation. **Alessandra Calcagno**: Writing – review & editing. **Marcello Massimini**: Writing – review & editing, Funding acquisition, Conceptualization. **Simone Sarasso**: Writing – review & editing, Conceptualization. **Mario Rosanova**: Writing – review & editing, Investigation, Data Curation, Conceptualization. **Silvia Casarotto**: Writing – review & editing, Writing – original draft, Visualization, Supervision, Project administration, Investigation, Data curation, Conceptualization.

## Acknowledgements

This work was financially supported by the following entities: ERC-2022-SYG Grant number 101071900 neurological mechanisms of injury and sleep-like cellular dynamics (NEMESIS); Italian National Recovery and Resilience Plan (NRRP), M4C2, funded by the European Union - NextGenerationEU (Project IR0000011, CUP B51E22000150006, “EBRAINS-Italy”); Canadian Institute for Advanced Research (CIFAR), Canada.

## References

Alexander, G.E., DeLong, M.R., Strick, P.L., 1986. Parallel organization of functionally segregated circuits linking basal ganglia and cortex. Annu Rev Neurosci 9, 357–381. 10.1146/annurev.ne.09.030186.002041

Andersen, R.A., Cui, H., 2009. Intention, Action Planning, and Decision Making in Parietal-Frontal Circuits. Neuron 63, 568–583. 10.1016/j.neuron.2009.08.028

Atalay, A.S., Fecchio, M., Edlow, B.L., 2025. Visualizing effective connectivity in the human brain. Brain Stimulation: Basic, Translational, and Clinical Research in Neuromodulation 18, 1254–1256. 10.1016/j.brs.2025.06.015

Beck, M.M., Heyl, M., Mejer, L., Vinding, M.C., Christiansen, L., Tomasevic, L., Siebner, H.R., 2024. Methodological Choices Matter: A Systematic Comparison of TMS-EEG Studies Targeting the Primary Motor Cortex. Human Brain Mapping 45, e70048. 10.1002/hbm.70048

Bestmann, S., Baudewig, J., Siebner, H.R., Rothwell, J.C., Frahm, J., 2004. Functional MRI of the immediate impact of transcranial magnetic stimulation on cortical and subcortical motor circuits. Eur J Neurosci 19, 1950–1962. 10.1111/j.1460-9568.2004.03277.x

Casali, A.G., Casarotto, S., Rosanova, M., Mariotti, M., Massimini, M., 2010. General indices to characterize the electrical response of the cerebral cortex to TMS. NeuroImage 49, 1459–1468. 10.1016/j.neuroimage.2009.09.026

Casarotto, S., Fecchio, M., Rosanova, M., Varone, G., D’Ambrosio, S., Sarasso, S., Pigorini, A., Russo, S., Comanducci, A., Ilmoniemi, R.J., Massimini, M., 2022. The rt-TEP tool: real-time visualization of TMS-Evoked Potentials to maximize cortical activation and minimize artifacts. Journal of Neuroscience Methods 370, 109486. 10.1016/j.jneumeth.2022.109486

Casarotto, S., Hassan, G., Rosanova, M., Sarasso, S., Derchi, C., Trimarchi, P.D., Viganò, A., Russo, S., Fecchio, M., Devalle, G., Navarro, J., Massimini, M., Comanducci, A., 2024. Dissociations between spontaneous electroencephalographic features and the perturbational complexity index in the minimally conscious state. Eur J of Neuroscience 59, 934–947. 10.1111/ejn.16299

Casarotto, S., Turco, F., Comanducci, A., Perretti, A., Marotta, G., Pezzoli, G., Rosanova, M., Isaias, I.U., 2019. Excitability of the supplementary motor area in Parkinson’s disease depends on subcortical damage. Brain Stimul 12, 152–160. 10.1016/j.brs.2018.10.011

Chiappa, K.H., Ropper, A.H., 1982. Evoked Potentials in Clinical Medicine. New England Journal of Medicine 306, 1140–1150. 10.1056/NEJM198205133061904

Cona, F., Zavaglia, M., Massimini, M., Rosanova, M., Ursino, M., 2011. A neural mass model of interconnected regions simulates rhythm propagation observed via TMS-EEG. NeuroImage, Special Issue: Educational Neuroscience 57, 1045–1058. 10.1016/j.neuroimage.2011.05.007

Cruccu, G., Aminoff, M.J., Curio, G., Guerit, J.M., Kakigi, R., Mauguiere, F., Rossini, P.M., Treede, R.-D., Garcia-Larrea, L., 2008. Recommendations for the clinical use of somatosensory-evoked potentials. Clinical Neurophysiology 119, 1705–1719. 10.1016/j.clinph.2008.03.016

Dawson, G.D., 1954. A summation technique for the detection of small evoked potentials. Electroencephalography and Clinical Neurophysiology 6, 65–84. 10.1016/0013-4694(54)90007-3

Denslow, S., Lomarev, M., George, M.S., Bohning, D.E., 2005. Cortical and subcortical brain effects of Transcranial Magnetic Stimulation (TMS)-induced movement: An interleaved TMS/functional magnetic resonance imaging study. Biological Psychiatry 57, 752–760. 10.1016/j.biopsych.2004.12.017

Destrieux, C., Fischl, B., Dale, A., Halgren, E., 2010. Automatic parcellation of human cortical gyri and sulci using standard anatomical nomenclature. NeuroImage 53, 1–15. 10.1016/j.neuroimage.2010.06.010

Donati, F.L., Mayeli, A., Sharma, K., Janssen, S.A., Lagoy, A.D., Casali, A.G., Ferrarelli, F., 2023. Natural Oscillatory Frequency Slowing in the Premotor Cortex of Early-Course Schizophrenia Patients: A TMS-EEG Study. Brain Sciences 13, 534. 10.3390/brainsci13040534

Farzan, F., Bortoletto, M., 2022. Identification and verification of a “true” TMS evoked potential in TMS-EEG. J Neurosci Methods 378, 109651. 10.1016/j.jneumeth.2022.109651

Fecchio, M., Pigorini, A., Comanducci, A., Sarasso, S., Casarotto, S., Premoli, I., Derchi, C.-C., Mazza, A., Russo, S., Resta, F., Ferrarelli, F., Mariotti, M., Ziemann, U., Massimini, M., Rosanova, M., 2017. The spectral features of EEG responses to transcranial magnetic stimulation of the primary motor cortex depend on the amplitude of the motor evoked potentials. PLOS ONE 12, e0184910. 10.1371/journal.pone.0184910

Ferrarelli, F., Massimini, M., Sarasso, S., Casali, A., Riedner, B.A., Angelini, G., Tononi, G., Pearce, R.A., 2010. Breakdown in cortical effective connectivity during midazolam-induced loss of consciousness. Proc Natl Acad Sci U S A 107, 2681–2686. 10.1073/pnas.0913008107

Ferrarelli, F., Sarasso, S., Guller, Y., Riedner, B.A., Peterson, M.J., Bellesi, M., Massimini, M., Postle, B.R., Tononi, G., 2012. Reduced Natural Oscillatory Frequency of Frontal Thalamocortical Circuits in Schizophrenia. Arch Gen Psychiatry 69. 10.1001/archgenpsychiatry.2012.147

Fischl, B., 2012. FreeSurfer. Neuroimage 62, 774–781. 10.1016/j.neuroimage.2012.01.021

Frauscher, B., von Ellenrieder, N., Zelmann, R., Doležalová, I., Minotti, L., Olivier, A., Hall, J., Hoffmann, D., Nguyen, D.K., Kahane, P., Dubeau, F., Gotman, J., 2018. Atlas of the normal intracranial electroencephalogram: neurophysiological awake activity in different cortical areas. Brain 141, 1130–1144. 10.1093/brain/awy035

Gilbert, C.D., Wiesel, T.N., 1989. Columnar specificity of intrinsic horizontal and corticocortical connections in cat visual cortex. J Neurosci 9, 2432–2442. 10.1523/JNEUROSCI.09-07-02432.1989

Halliday, A.M., McDonald, W.I., Mushin, J., 1972. Delayed visual evoked response in optic neuritis. Lancet 1, 982–985. 10.1016/s0140-6736(72)91155-5

Janssen, A.M., Oostendorp, T.F., Stegeman, D.F., 2015. The coil orientation dependency of the electric field induced by TMS for M1 and other brain areas. J Neuroeng Rehabil 12, 47. 10.1186/s12984-015-0036-2

Jones, E.G., 2001. The thalamic matrix and thalamocortical synchrony. Trends in Neurosciences 24, 595–601. 10.1016/S0166-2236(00)01922-6

Logi, F., Fischer, C., Murri, L., Mauguière, F., 2003. The prognostic value of evoked responses from primary somatosensory and auditory cortex in comatose patients. Clinical Neurophysiology 114, 1615–1627. 10.1016/S1388-2457(03)00086-5

Lőrincz, M.L., Kékesi, K.A., Juhász, G., Crunelli, V., Hughes, S.W., 2009. Temporal Framing of Thalamic Relay-Mode Firing by Phasic Inhibition during the Alpha Rhythm. Neuron 63, 683–696. 10.1016/j.neuron.2009.08.012

Lysomiski, A.L., Dijk, J.V., Giampiccolo, D., 2026. Mapping direct cortical responses to their underlying cytoarchitectonics. Clinical Neurophysiology 184, 2111505. 10.1016/j.clinph.2026.2111505

Markov, N.T., Ercsey-Ravasz, M.M., Ribeiro Gomes, A.R., Lamy, C., Magrou, L., Vezoli, J., Misery, P., Falchier, A., Quilodran, R., Gariel, M.A., Sallet, J., Gamanut, R., Huissoud, C., Clavagnier, S., Giroud, P., Sappey-Marinier, D., Barone, P., Dehay, C., Toroczkai, Z., Knoblauch, K., Van Essen, D.C., Kennedy, H., 2014. A Weighted and Directed Interareal Connectivity Matrix for Macaque Cerebral Cortex. Cereb Cortex 24, 17–36. 10.1093/cercor/bhs270

Massimini, M., Ferrarelli, F., Huber, R., Esser, S.K., Singh, H., Tononi, G., 2005. Breakdown of cortical effective connectivity during sleep. Science 309, 2228–2232. 10.1126/science.1117256

McHaffie, J.G., Stanford, T.R., Stein, B.E., Coizet, V., Redgrave, P., 2005. Subcortical loops through the basal ganglia. Trends in Neurosciences 28, 401–407. 10.1016/j.tins.2005.06.006

Meneghini, G., Engelhardt, M., Burzlaff, M., Zaykova, A., Vajkoczy, P., Lioumis, P., Rosanova, M., Picht, T., 2025. Probing cortical reactivity before and after brain tumor resection: A TMS-EEG case. Brain Stimulation: Basic, Translational, and Clinical Research in Neuromodulation 18, 19–21. 10.1016/j.brs.2024.12.004

Moliadze, V., Zhao, Y., Eysel, U., Funke, K., 2003. Effect of transcranial magnetic stimulation on single-unit activity in the cat primary visual cortex. The Journal of Physiology 553, 665–679. 10.1113/jphysiol.2003.050153

Momi, D., Wang, Z., Parmigiani, S., Mikulan, E., Bastiaens, S.P., Oveisi, M.P., Kadak, K., Gaglioti, G., Waters, A.C., Hill, S., Pigorini, A., Keller, C.J., Griffiths, J.D., 2025. Stimulation mapping and whole-brain modeling reveal gradients of excitability and recurrence in cortical networks. Nat Commun 16, 3222. 10.1038/s41467-025-58187-6

Mueller, J.K., Grigsby, E.M., Prevosto, V., Petraglia, F.W., Rao, H., Deng, Z.-D., Peterchev, A.V., Sommer, M.A., Egner, T., Platt, M.L., Grill, W.M., 2014. Simultaneous transcranial magnetic stimulation and single-neuron recording in alert non-human primates. Nat Neurosci 17, 1130–1136. 10.1038/nn.3751

Opitz, A., Windhoff, M., Heidemann, R.M., Turner, R., Thielscher, A., 2011. How the brain tissue shapes the electric field induced by transcranial magnetic stimulation. Neuroimage 58, 849–859. 10.1016/j.neuroimage.2011.06.069

Parmigiani, S., Mikulan, E., Russo, S., Sarasso, S., Zauli, F.M., Rubino, A., Cattani, A., Fecchio, M., Giampiccolo, D., Lanzone, J., D’Orio, P., Del Vecchio, M., Avanzini, P., Nobili, L., Sartori, I., Massimini, M., Pigorini, A., 2022. Simultaneous stereo-EEG and high-density scalp EEG recordings to study the effects of intracerebral stimulation parameters. Brain Stimulation 15, 664–675. 10.1016/j.brs.2022.04.007

Peters, J.C., Reithler, J., Graaf, T.A. de, Schuhmann, T., Goebel, R., Sack, A.T., 2020. Concurrent human TMS-EEG-fMRI enables monitoring of oscillatory brain state-dependent gating of cortico-subcortical network activity. Commun Biol 3, 40. 10.1038/s42003-020-0764-0

Pigorini, A., Sarasso, S., Proserpio, P., Szymanski, C., Arnulfo, G., Casarotto, S., Fecchio, M., Rosanova, M., Mariotti, M., Lo Russo, G., Palva, J.M., Nobili, L., Massimini, M., 2015. Bistability breaks-off deterministic responses to intracortical stimulation during non-REM sleep. NeuroImage 112, 105–113. 10.1016/j.neuroimage.2015.02.056

Premoli, I., Rivolta, D., Espenhahn, S., Castellanos, N., Belardinelli, P., Ziemann, U., Müller-Dahlhaus, F., 2014. Characterization of GABAB-receptor mediated neurotransmission in the human cortex by paired-pulse TMS–EEG. NeuroImage 103, 152–162. 10.1016/j.neuroimage.2014.09.028

Reinhold, K., Lien, A.D., Scanziani, M., 2015. Distinct recurrent versus afferent dynamics in cortical visual processing. Nat Neurosci 18, 1789–1797. 10.1038/nn.4153

Rolls, E.T., Deco, G., Huang, C.-C., Feng, J., 2022. The human posterior parietal cortex: effective connectome, and its relation to function. Cereb Cortex 33, 3142–3170. 10.1093/cercor/bhac266

Romei, V., Brodbeck, V., Michel, C., Amedi, A., Pascual-Leone, A., Thut, G., 2008. Spontaneous fluctuations in posterior alpha-band EEG activity reflect variability in excitability of human visual areas. Cereb Cortex 18, 2010–2018. 10.1093/cercor/bhm229

Rosanova, M., Casali, A., Bellina, V., Resta, F., Mariotti, M., Massimini, M., 2009. Natural Frequencies of Human Corticothalamic Circuits. Journal of Neuroscience 29, 7679–7685. 10.1523/JNEUROSCI.0445-09.2009

Rosanova, M., Fecchio, M., Casarotto, S., Sarasso, S., Casali, A.G., Pigorini, A., Comanducci, A., Seregni, F., Devalle, G., Citerio, G., Bodart, O., Boly, M., Gosseries, O., Laureys, S., Massimini, M., 2018. Sleep-like cortical OFF-periods disrupt causality and complexity in the brain of unresponsive wakefulness syndrome patients. Nat Commun 9, 4427. 10.1038/s41467-018-06871-1

Russo, S., Claar, L.D., Furregoni, G., Marks, L.C., Krishnan, G., Zauli, F.M., Hassan, G., Solbiati, M., d’Orio, P., Mikulan, E., Sarasso, S., Rosanova, M., Sartori, I., Bazhenov, M., Pigorini, A., Massimini, M., Koch, C., Rembado, I., 2025. Thalamic feedback shapes brain responses evoked by cortical stimulation in mice and humans. Nat Commun 16, 3627. 10.1038/s41467-025-58717-2

Russo, S., Sarasso, S., Puglisi, G.E., Dal Palù, D., Pigorini, A., Casarotto, S., D’Ambrosio, S., Astolfi, A., Massimini, M., Rosanova, M., Fecchio, M., 2022. TAAC - TMS Adaptable Auditory Control: A universal tool to mask TMS clicks. Journal of Neuroscience Methods 370, 109491. 10.1016/j.jneumeth.2022.109491

Sarasso, S., D’Ambrosio, S., Fecchio, M., Casarotto, S., Viganò, A., Landi, C., Mattavelli, G., Gosseries, O., Quarenghi, M., Laureys, S., Devalle, G., Rosanova, M., Massimini, M., 2020. Local sleep-like cortical reactivity in the awake brain after focal injury. Brain 143, 3672–3684. 10.1093/brain/awaa338

Sarasso, S., Rosanova, M., Casali, A.G., Casarotto, S., Fecchio, M., Boly, M., Gosseries, O., Tononi, G., Laureys, S., Massimini, M., 2014. Quantifying Cortical EEG Responses to TMS in (Un)consciousness. Clin EEG Neurosci 45, 40–49. 10.1177/1550059413513723

Solomon, E.A., Wang, J.B., Oya, H., Howard, M.A., Trapp, N.T., Uitermarkt, B.D., Boes, A.D., Keller, C.J., 2024. TMS provokes target-dependent intracranial rhythms across human cortical and subcortical sites. Brain Stimul 17, 698–712. 10.1016/j.brs.2024.05.014

Starr, A., Achor, L.J., 1975. Auditory Brain Stem Responses in Neurological Disease. Arch Neurol 32, 761–768. 10.1001/archneur.1975.00490530083009

Stettler, D.D., Das, A., Bennett, J., Gilbert, C.D., 2002. Lateral Connectivity and Contextual Interactions in Macaque Primary Visual Cortex. Neuron 36, 739–750. 10.1016/S0896-6273(02)01029-2

Stockwell, R.G., Mansinha, L., Lowe, R.P., 1996. Localization of the complex spectrum: the S transform. IEEE Trans. Signal Process. 44, 998–1001. 10.1109/78.492555

Thomas Yeo, B.T., Krienen, F.M., Sepulcre, J., Sabuncu, M.R., Lashkari, D., Hollinshead, M., Roffman, J.L., Smoller, J.W., Zöllei, L., Polimeni, J.R., Fischl, B., Liu, H., Buckner, R.L., 2011. The organization of the human cerebral cortex estimated by intrinsic functional connectivity. J Neurophysiol 106, 1125–1165. 10.1152/jn.00338.2011

Ts’o, D.Y., Frostig, R.D., Lieke, E.E., Grinvald, A., 1990. Functional Organization of Primate Visual Cortex Revealed by High Resolution Optical Imaging. Science 249, 417–420. 10.1126/science.2165630

van den Heuvel, M.P., Sporns, O., 2013. Network hubs in the human brain. Trends Cogn Sci 17, 683–696. 10.1016/j.tics.2013.09.012

Ziemann, U., Bai, Y., Baumer, F.M., Beck, M.M., Belardinelli, P., Belvisi, D., Bender, S., Bergmann, T.O., Bortoletto, M., Casarotto, S., Casula, E., Chaves, A.R., De Andrade, D.C., Conte, A., Daskalakis, Z.J., Farzan, F., Ferrarelli, F., Fitzgerald, P.B., Gordon, P.C., Grefkes, C., Harquel, S., Hernandez-Pavon, J.C., Hill, A.T., Hoy, K.E., Hummel, F.C., Julkunen, P., Kallioniemi, E., Keller, C.J., Kimiskidis, V.K., Kirkovski, M., Koch, G., Leodori, G., Lioumis, P., Määttä, S., Maidan, I., Massimini, M., Mengel, A., Metsomaa, J., Miniussi, C., Mutanen, T.P., Noda, Y., Ozdemir, R.A., Raffin, E., Rocchi, L., Rogasch, N.C., Rosanova, M., Santarnecchi, E., Sarasso, S., Schabrun, S.M., Shafi, M.M., Siebner, H.R., Tolner, E.A., Tomasevic, L., Tremblay, S., Tscherpel, C., Veniero, D., Versace, V., Voineskos, D., Vucic, S., Zangen, A., Zrenner, C., Ilmoniemi, R.J., 2026. Clinical utility and prospective of TMS–EEG: Updated review from an international expert group. Clinical Neurophysiology 184, 2111487. 10.1016/j.clinph.2025.2111487

