## Supplementary materials for "Temporal fingerprints of TMS-evoked potentials across thalamocortical circuits"

### Supplementary Material

#### Supplementary Methods

##### Instrumentation

Single, biphasic TMS pulses were administered with a focal figure-of-8 coil (mean/outer winding diameter ca. 50/70 mm, pulse length ca. 280 µs, focal area of the stimulation hot spot .68 $cm^2$). Stimulation parameters were precisely controlled through a Navigated Brain Stimulation (NBS) system integrated within the main TMS unit (Nexstim Ltd., Finland). Cortical targets were selected on individual T1-weighted magnetic resonance images (MRI). For each target, the 3D spatial coordinates of the maximum of the induced electric field (EF-max) on the cortex as well as its intensity in V/m were automatically stored in a logfile.
EEG responses to TMS were recorded with a TMS-compatible Brainamp DC amplifier (Brain Products GmbH, Germany) connected to a 64-channel EEG cap with C-ring shape, Ag/AgCl electrodes located according to the 10-10 montage. Two channels were used for recording electrooculographic (EOG) activity with a diagonal montage. Data were collected with a hardware filtering bandwidth between DC and 1000 Hz and were sampled at 5000 Hz with 0.5µV amplitude resolution. Additional reference and ground electrodes were placed on the forehead over frontal sinuses. Impedance was kept below 5 kΩ in all channels.

##### Data pre-processing

The artifact induced by the TMS coil discharge was removed from raw recordings by replacing the EEG signal from −2 to +5 ms around each pulse with a baseline segment (mirrored from -2 to -9 ms and sign-flipped) and by subsequently applying a 4-ms long moving-average filter. Continuous EEG data were detrended by high-pass filtering at .01 Hz with a 1st order IIR filter, high-pass filtered at 1 Hz using a zero-phase Butterworth filter (3rd order) and then segmented into trials spanning ±600 ms around TMS pulses. Single trials and channels visually affected by sporadic artifacts were rejected from further analysis. Then, single-trial TEPs were re-referenced to the average reference and baseline-corrected. Independent Component Analysis (ICA), performed with the *runica* function implemented in EEGLAB (Delorme & Makeig, 2004), was used to identify and remove components associated with ocular and muscular artifacts. Reconstructed TEPs were low-pass filtered at 45 Hz (zero-phase Butterworth filter, 3rd order) and downsampled to 1000 Hz. Finally, rejected channels were interpolated using spherical splines.

### Supplementary Figures

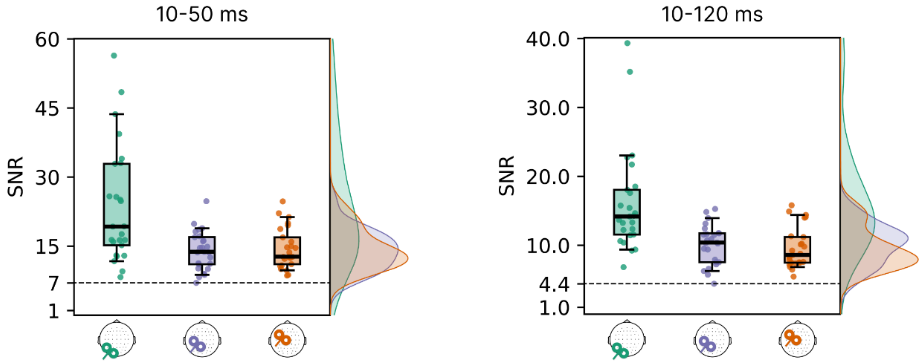


**Figure S1. Signal-to-noise ratio across targets.** Boxplots show the distribution across sessions of the average SNR of the four EEG channels selected for each subject (see Section 3.1 in the main results), computed as the ratio between the mean absolute amplitude in a post-stimulus window (10–50 ms, left; 10–120 ms, right) and the mean absolute amplitude in the pre-stimulus window (−400 to −10 ms). The dashed horizontal lines indicate the minimum SNR value observed across all sessions, corresponding to 7 (10–50 ms) and 4.4 (10–120 ms) in the respective time windows.

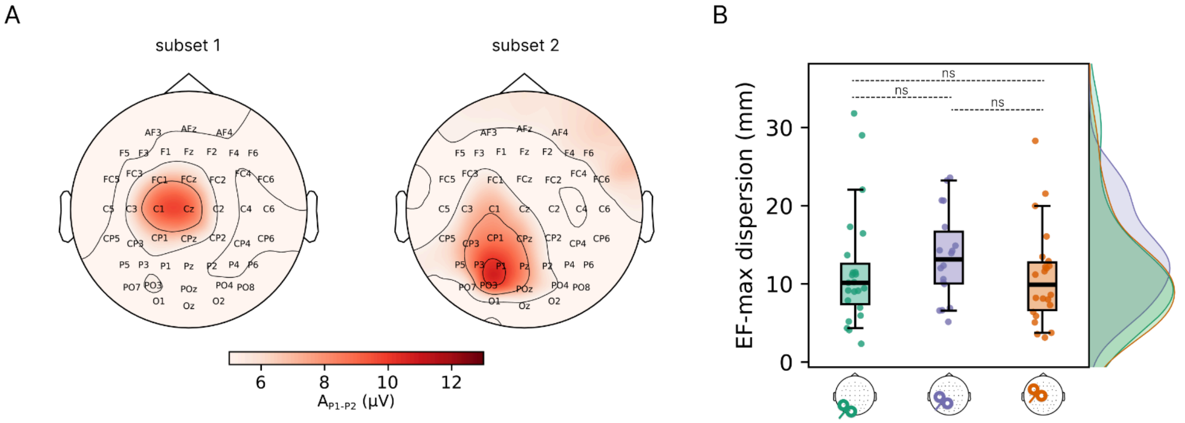


**Figure S2. Spatial distribution of TEPs amplitude and of anatomical targets. (A)** Topographic maps of $A_{P1-P2}$ of parietal TEPs averaged across 12 subjects (subset 1, left) and 11 subjects (subset 2, right). Subsets 1 and 2 were created by grouping subjects who showed either C1 or P1, respectively, among the individual ROI channels automatically identified from parietal TEPs with *ntop* = 4. These topographic maps show a similar focality to the ones obtained for occipital and frontal TEPs (Figure 2A main text). **(B)** Euclidean distance between single-subject EF-max locations for each TMS target and the corresponding centroid (EF-max dispersion). Pairwise differences between targets were assessed using two-sided Mann–Whitney U tests (all comparisons not significant, $p \geq 0.05$; ns).

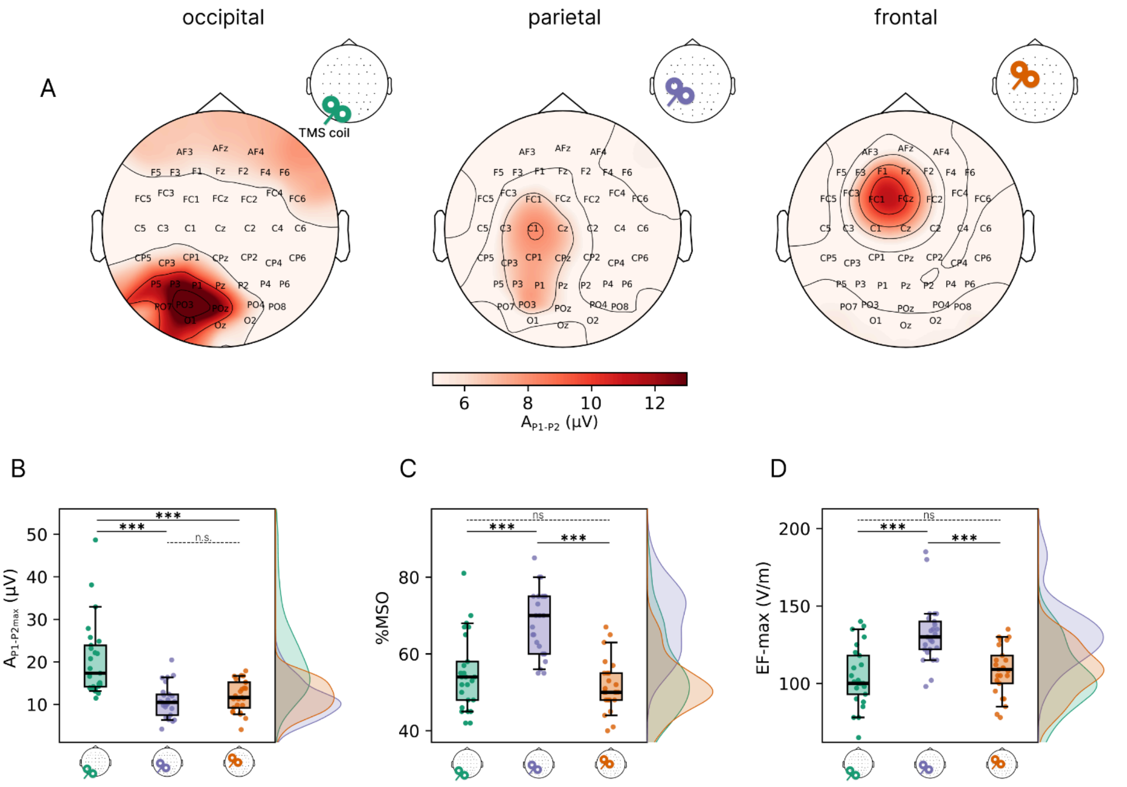


**Figure S3. Comparison of TEPs amplitude and stimulation intensity across targets. (A)** Grand-average topographic maps of $A_{P1\text{-}P2}$ for occipital, parietal, and frontal stimulation sites. **(B)** Maximum $A_{P1\text{-}P2}$ value across channels ($A_{P1\text{-}P2max}$) in occipital (green), parietal (violet) and frontal (orange) TEPs. **(C-D)** Stimulation intensity, expressed as percentage of the maximal stimulator output (%MSO, C) and expressed as maximum value of the induced electric field (EF-max, D), applied in occipital (green), parietal (violet) and frontal (orange) targets. In all panels, colored dots represent individual values whereas group-level distributions in each cortical target are depicted as boxplots (median, 25th and 75th percentile, and 5th and 95th percentile) and as density plots. P-values were obtained by applying the Kruskal-Wallis test separately to each measure, followed by pairwise Dunn's post-hoc tests with Holm-Bonferroni correction (where significance is indicated as * for $p < 0.05$, ** for $p < 0.01$, *** for $p < 0.001$, and ns for non-significant comparisons ($p \geq 0.05$).

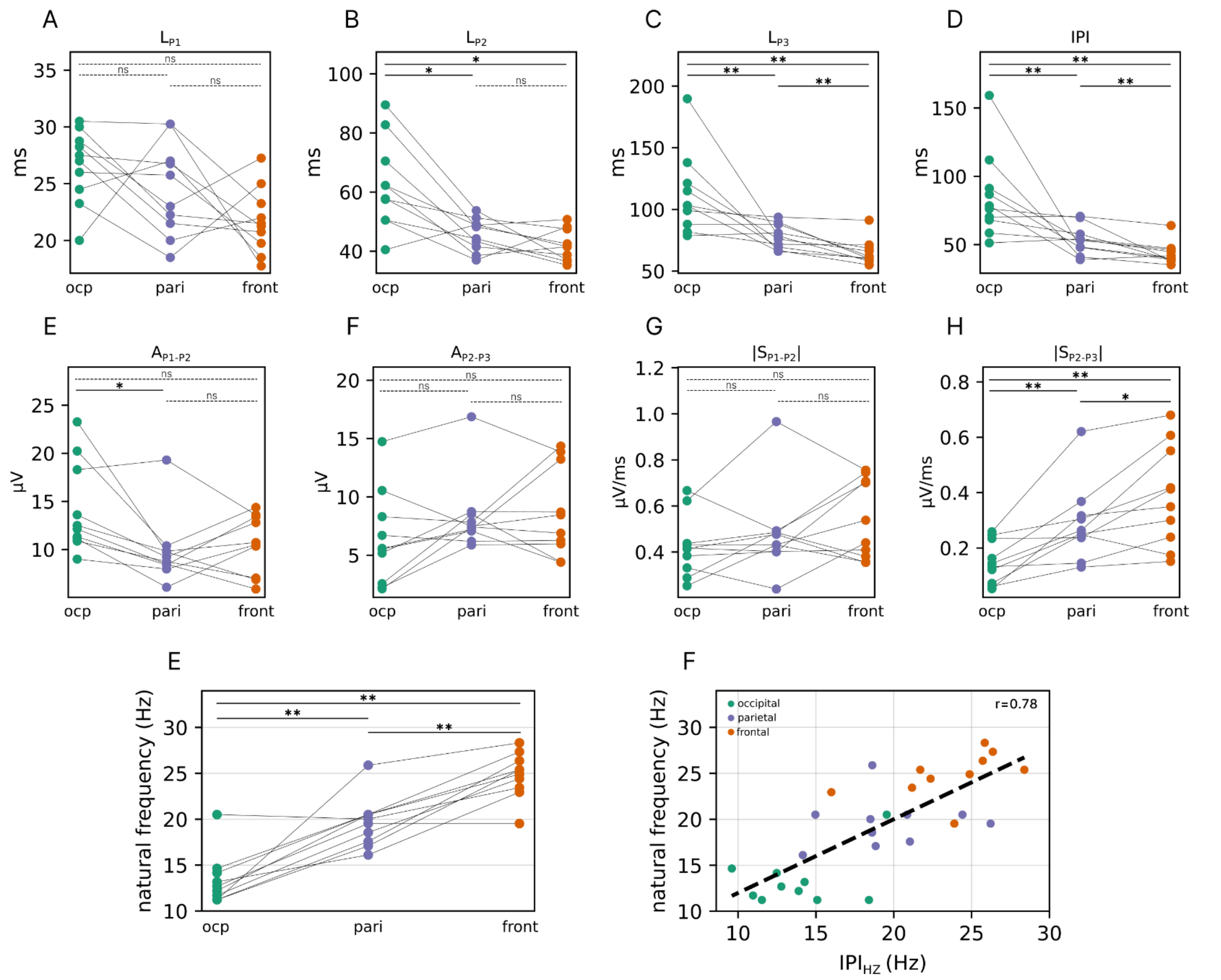


**Figure S4. Within-subject comparisons of morphological and spectral features of TEPs across stimulation sites. (A–H)** Group-level distributions of LP1 (A), LP2 (B), LP3 (C), IPI (D), AP1-P2 (E), AP2-P3 (F), |SP1-P2| (G) and |SP2-P3| (H) are depicted as boxplots (median, 25th and 75th percentile and 5th and 95th) for occipital (green), parietal (violet) and frontal (orange) targets. These measures represent a subset of the ones reported in Figure 2 in the main text because they were obtained from the 10 subjects who were stimulated in all the three cortical sites (see Methods). Reported P-values were obtained by applying the Kruskal-Wallis test separately to each measure, followed by Dunn's pairwise post-hoc tests with Holm-Bonferroni correction. **(E)** Natural frequency estimated from time–frequency analysis for each site, shown for each participant (colored points) who received all three stimulations. In the panels, each line connects measurements from the same subject. Within-subject analyses for participants receiving all three stimulations ($n = 10$) were performed using Friedman tests with post-hoc Wilcoxon signed-rank tests and Holm–Bonferroni correction; p-values are shown in the figure and significance is indicated as * for $p < 0.05$, ** for $p < 0.01$, *** for $p < 0.001$, and ns for non-significant comparisons ($p \geq 0.05$). **(F)** Correlation between natural frequency and the reciprocal of the interpeak time (1/interpeak) in the paired subgroup, confirming a strong relationship ($r = 0.78$, $p = 4\times10^{-7}$) between these temporal and spectral measures.

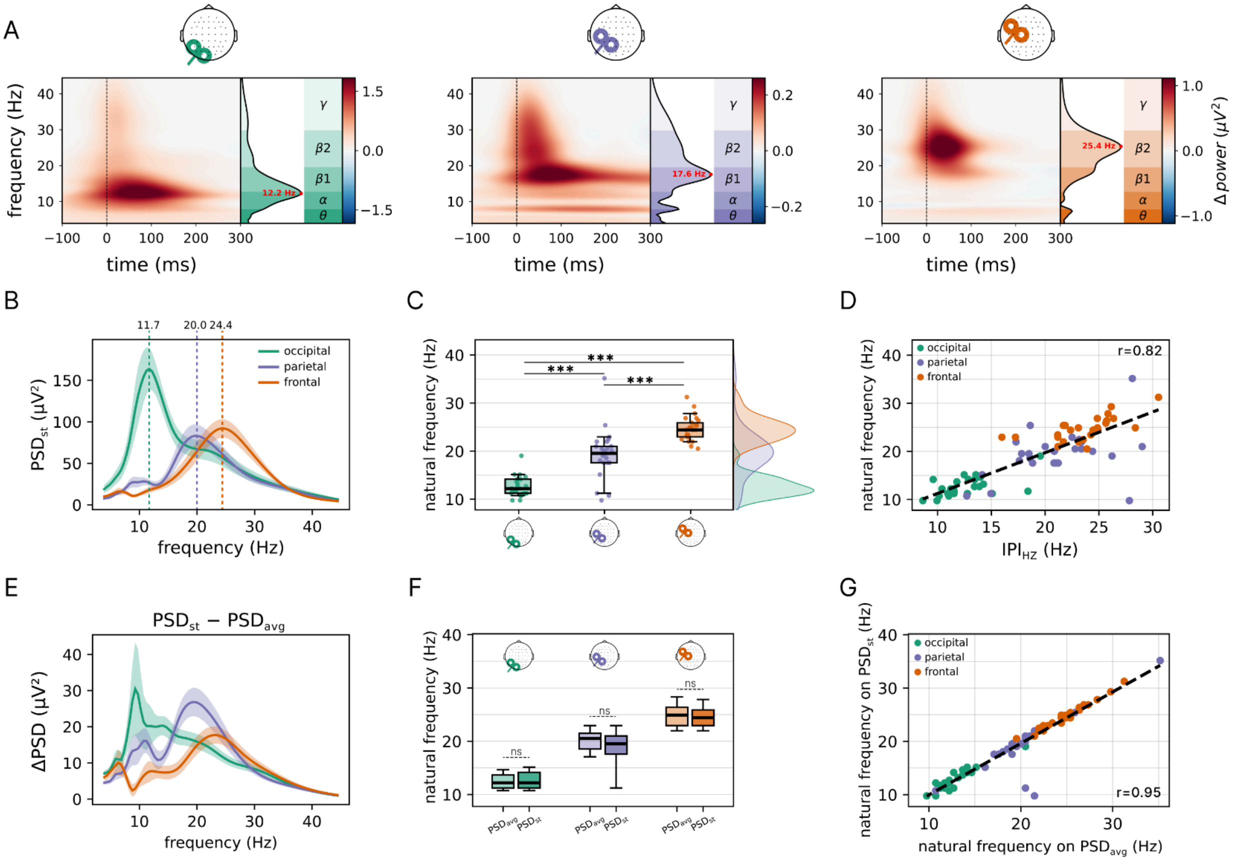


**Figure S5. Single-trial time–frequency analysis reproduces stimulation-specific natural frequencies. (A)** Time–frequency power representations of the TEP for a representative subject (as in Figure 4 in the main text) following occipital (left), parietal (middle), and frontal (right) stimulation, computed using the Stockwell transform on the four automatically selected channels. Time–frequency power was computed at the single-trial level and then averaged across trials ($TF_{st}$), averaged across the four channels, and baseline-corrected by subtracting the average pre-stimulus power within each frequency bin. The single-trial power spectrum ($PSD_{st}$) was then obtained by summing $TF_{st}$ power over the post-stimulus window 20–120 ms (spectral plots shown to the right of each panel), and the frequency corresponding to the peak of the evoked power was defined as the natural frequency (red dot; value labeled). **(B)** Grand-average $PSD_{st}$ across subjects for the three stimulation conditions (shaded areas indicate SEM). **(C)** Box plots of the natural frequency derived from $PSD_{st}$ across subjects for each stimulation site (occipital: green, parietal: violet, frontal:: orange); post-hoc pairwise comparisons (Dunn test) are Holm–Bonferroni corrected (significance reported in the panel as in Figure 4). Density plots to the right summarize the natural frequency distribution across subjects for each stimulation site. **(D)** Positive linear association between natural frequency ($PSD_{st}$) and $IPI_{Hz}$ ($r = 0.82$; $p = 3\times10^{-19}$); each point represents one subject and stimulation site. **(E)** Grand-average difference spectrum between $PSD_{avg}$ (Figure 4) and $PSD_{st}$, showing that residual post-stimulus power (ΔPSD) preserves a frequency gradient across stimulation targets. **(F)** Within-target comparison of natural frequency obtained from $PSD_{avg}$ versus $PSD_{st}$ shows no significant differences (ns). **(G)** Strong linear relationship between natural frequency computed on PSDavg and PSDst ($r = 0.95$; $p = 4\times10^{-40}$)

#### Robustness of the correlation between spectral and temporal features across analysis parameters

The results reported in figure 4, S4 (E-F) and S5 were obtained using a specific set of analysis parameters: ntop = 4 ROI channels, a Gaussian window width of 0.7 for the Stockwell transform, and a post-stimulus time window of 20-120 ms for computing the evoked PSD. To assess the stability of the main findings and to cement the relation between temporal and spectral features, we systematically varied each of these parameters and tested whether the NF-$IPI_{Hz}$ correlation and the between-site differences in NF were preserved across the resulting combinations (Fig. S6). NF and $IPI_{Hz}$ remained highly correlated across a wide range of parameter settings, with the peak correlation tending to shift towards larger width values as the power cumulation window was extended. Significant differences in NF between all pairs of stimulation sites were consistently observed across the full range of widths, ntop values, and time windows tested, confirming that the posterior-to-anterior spectral gradient is a robust feature of the data and not a product of any particular analysis choice.

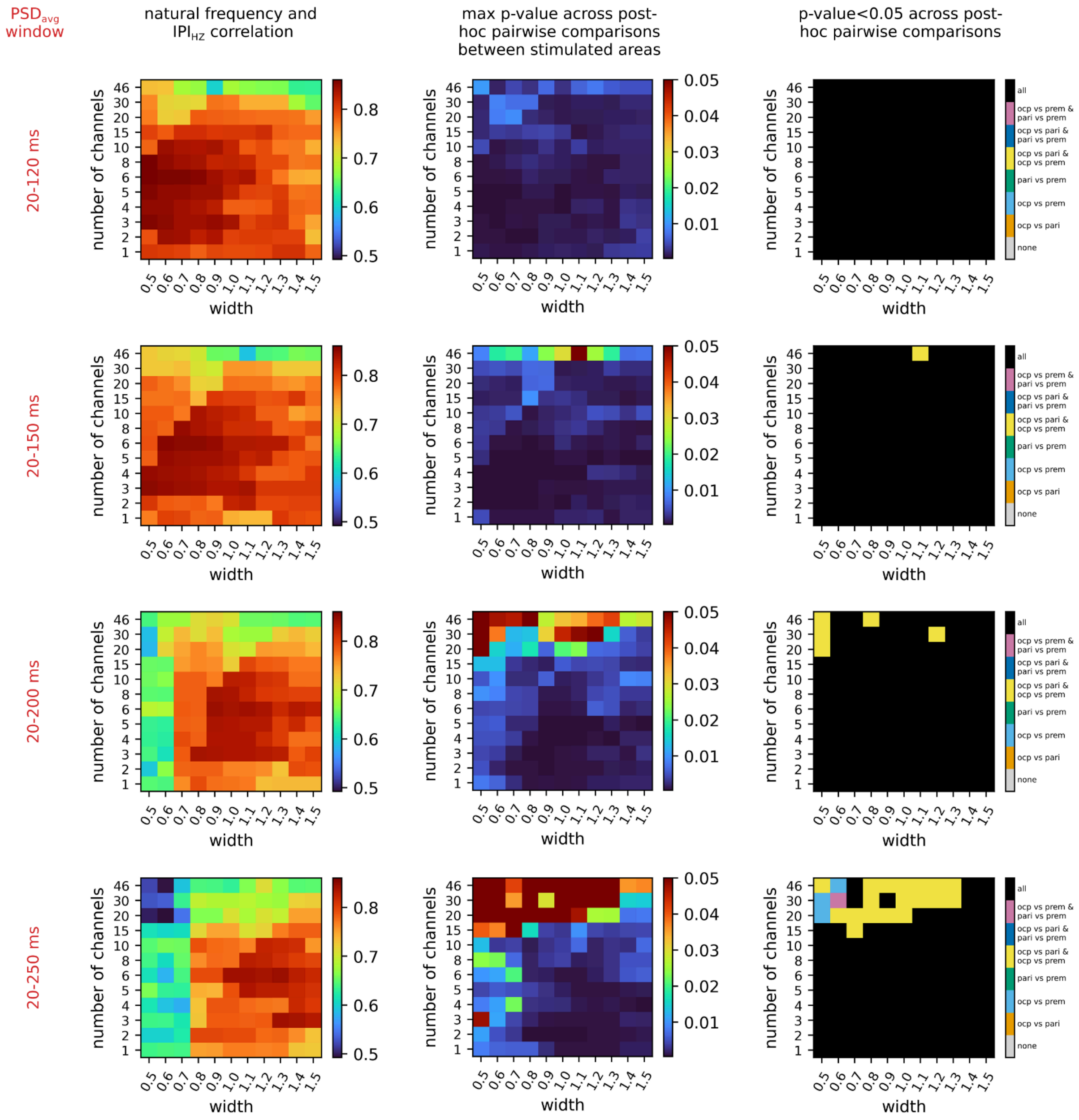


**Figure S6. Robustness of natural frequency estimation across parameter variations.** Parameter exploration analysis demonstrating stability of natural frequency metrics. Each row represents a different $PSD_{avg}$ summation window: 20–120 ms (top), 20–150 ms, 20–200 ms, and 20–250 ms (bottom). Columns 1-3 display heatmaps evaluating the effects of Stockwell transform width (x-axis) and number of selected channels (y-axis; via *ntop* parameter) on key statistical metrics. Column 1: Correlation coefficients (r) between natural frequency and 1/interpeak time. Red hues indicate strong correlations ($r > 0.8$), confirming the robustness of this relationship across parameter combinations. Column 2: Maximum p-values across all post-hoc pairwise comparisons (occipital vs. parietal, occipital vs. frontal, parietal vs. frontal). Dark red indicates parameter combinations where at least one comparison was non-significant ($p > 0.05$). Column 3: Significance patterns of post-hoc comparisons. Colors denote specific combinations of significant pairwise differences (see colorbar), with black indicating all three comparisons remained significant ($p < 0.05$). P-values are corrected with the Holm-Bonferroni method.
